# Adaptive immunity is required for durable responses to alectinib in murine models of EML4-ALK lung cancer

**DOI:** 10.1101/2022.04.14.488385

**Authors:** Emily K. Kleczko, Trista K. Hinz, Teresa T. Nguyen, Natalia J. Gurule, Andre Navarro, Anh T. Le, Amber M. Johnson, Jeff Kwak, Diana I. Polhac, Eric T. Clambey, Mary Weiser-Evans, Tejas Patil, Erin L. Schenk, Lynn E. Heasley, Raphael A. Nemenoff

## Abstract

**Purpose:** Lung cancers bearing oncogenic EML4-ALK fusions respond to targeted tyrosine kinase inhibitors (TKIs; e.g. alectinib), with variation in the degree of shrinkage and duration of treatment (DOT). We previously demonstrated a positive association of a TKI-induced interferon gamma (IFNγ) transcriptional response with DOT in EGFR-mutant lung cancers. Herein, we used murine models of EML4-ALK lung cancer to test a role for host immunity in the therapeutic response to alectinib.

**Experimental Design:** Three murine EML4-ALK cell lines (EA1, EA2, EA3) were implanted orthotopically into the lungs of immunocompetent and immunodeficient mice and treated with alectinib. Tumor volumes were serially measured by μCT. Immune cell content was measured by flow cytometry, multispectral immunofluorescence and CyTOF. Transcriptional responses to alectinib were assessed by RNAseq and secreted chemokines were measured by ELISA.

**Results:** All cell lines were sensitive to alectinib *in vitro*. EA1 and EA3 tumors retained residual disease that rapidly progressed upon termination of treatment while EA2 tumors were eliminated by TKI treatment. Alectinib induced inflammatory transcriptional programs and multiple chemokines in all cell lines while untreated tumors exhibited distinct baseline chemokine expression patterns and content of CD8+ T cells and myeloid subsets. When propagated in immune-deficient mice, all three cell line-derived lung tumor models exhibited significant shrinkage followed by prompt progression despite continuous alectinib treatment.

**Conclusions:** The findings support an hypothesis that host and TKI-stimulated production of chemokines by tumor cells promotes functional engagement of adaptive immune cells within the tumor microenvironment that enhances the durability and depth of TKI response.

**Statement of Translational Relevance:** Patients with metastatic lung cancer harboring ALK fusions are treated with targeted tyrosine kinase inhibitors (TKI) in the first line setting. Despite bearing the same driver oncogene, patients experience a range of tumor burden reduction and variable amounts of residual disease. Residual disease burden associates with patient survival and contributes to the emergence of drug resistance yielding treatment failure. The factors mediating this differential response to TKI and residual disease are incompletely understood. Our group has developed a panel of murine ALK driven lung cancer cell lines that reproducibly show differences in the depth and duration of response when implanted into immunocompetent mice. Data using this model indicate that the presence of CD8+ T cells is a major contributor to the depth and duration of response. These models will be critical in developing rational combination therapies to augment the immune microenvironment engagement along with TKIs to improve outcomes for these patients.

## Introduction

Lung adenocarcinomas driven by oncogenic tyrosine kinases including mutant EGFR and ALK fusions exhibit frequent and extensive responses to precision targeted tyrosine kinase inhibitors (TKIs), although with a wide range in the depth of response (DepOR) and time to progression or duration of treatment (DOT) (1-3). Alectinib, the current 1^st^ line standard of care for metastatic ALK+ lung cancers, yields objective tumor responses (<30% tumor shrinkage by RECIST) in ∼50-75% of TKI-treated patients and progression free survival (PFS) ranges from ∼10-30 months. Despite these successes, TKIs fail to completely eliminate tumor cells, with remaining cancer cells referred to as “drug tolerant persisters” (4) or “residual disease” (5). A clear association exists between initial DepOR and progression free and overall survival in ALK+ patients (6). Thus, understanding the biological underpinnings for variation in DepOR and time to progression may inform rational strategies for enhancing the efficacy of oncogene-targeted agents through novel drug combinations. Our recent published studies demonstrate that EGFR-targeted inhibitors induce an interferon (IFN) response program that varies markedly between distinct EGFR mutant lung cancer cell lines and positively associates with the duration of therapeutic response in EGFR-mutant lung cancer patients (7). In fact, there is a growing literature supporting the role of host immune cells in overall therapeutic response to precision oncology agents and cytotoxic drugs (8-11). However, the mechanisms whereby TKIs induce factors mediating paracrine signaling to the immune microenvironment, and the contribution to the overall therapeutic response are not well understood. This is critical to understanding how these pathways may be targeted for therapeutic gain.

While human samples can be used to develop correlations, preclinical mouse models that recapitulate critical features of the human disease open avenues to deep mechanistic exploration. To define the role of the tumor microenvironment (TME) in mediating response to TKI therapy in ALK positive lung cancer, we have used an orthotopic immunocompetent model whereby murine EML4-ALK fusion-positive lung cancer cells are directly implanted into the lungs of syngeneic C57BL/6 mice (12-14). In contrast to studies that have implanted lung cancer cell lines subcutaneously, this model requires tumors to develop in the relevant microenvironment of the lung such that the role of the adaptive and innate immune systems in contributing to the overall therapeutic response can be assessed. By examining a panel of cell lines with the same oncogenic driver, we sought to model the heterogeneity of response observed in patients. In this study, we demonstrate that distinct ALK fusion-bearing lung cancer cells exhibit differences in the depth and duration of response to the ALK inhibitor, alectinib. Furthermore, our data support a model in which differential interactions with the tumor microenvironment contribute to the depth of therapeutic response.

## Materials and Methods

### Establishment and culture of murine EML4-ALK cell lines

EA1 and EA3 cells were developed at the University of Colorado Anschutz Medical Center and EA1 cells have previously been described by our group (15,16). The EML4-ALK cell line referred to herein as EA2 was generously provided by Dr. Andrea Ventura at Memorial Sloan Kettering Cancer Center (17). To induce primary EML4-ALK tumors, a preparation of recombinant adenovirus encoding Cas9 and gRNAs targeting *Eml4* and *Alk* was purchased from ViraQuest Inc (North Liberty, IA). Tracheal administration of 50 μL Adeno-Cas9-gRNA virus at a dose of 1.5 × 10^8^ PFU/mouse induced multifocal tumor formation in both lungs after 8-12 weeks. Tumors were harvested, minced, and grown in Roswell Park Memorial Institute-1640 (RPMI, Corning) growth media supplemented with 10% fetal bovine serum (FBS, GIBCO) and 1% penicillin-streptomycin (Corning). Cells were maintained in culture and passaged until stable epithelial cell lines were established. Cells were cultured in a humidified incubator with 5% CO_2_ at 37°C.

### PCR Analysis to Confirm EML4-ALK Expression

To confirm the EML4-ALK inversion and fusion in the cell lines, RNA was isolated using Quick RNA Miniprep kit following manufacturer’s protocol (Zymo Research) and 5 μg of RNA was reverse transcribed using Maxima cDNA Synthesis Kit following manufacturer’s protocol (Fisher Scientific). The resulting cDNA was submitted to PCR analysis using PCRBIO VeriFi™ Mix (PCRBIOSYSTEMS) and previously reported oligonucleotide primers (Eml4-forward: 5’-GAGCCTTGTTGATACATCGTTC-3’ and Alk-reverse: 5’-CAAGGCAGTGAGAACCTGAA-3’ (17)). The PCR products (190 bp) were electrophoresed on 2% agarose gel, stained with ethidium bromide, and photographed.

### Immunoblotting

Cells were collected in phosphate-buffered saline, centrifuged, and suspended in lysis buffer (0.5% Triton X-100, 50 mM β-glycerophosphate (pH 7.2), 0.1 mM Na_3_VO_4_, 2 mM MgCl_2_, 1 mM EGTA, 1 mM DTT, 0.3 M NaCl, 2 μg/ml leupeptin and 4 μg/ml aprotinin). Aliquots of the cell lysates containing 50 μg of protein were submitted to SDS-PAGE and immunoblotted for TP53 (Cell Signaling Technology, #32532) and β-actin (Cell Signaling Technology, #4967) as a loading control.

### Enzyme-linked immunosorbent assay (ELISA)

Cells were seeded in 10-cm dishes and 24 hrs later, the cells were treated with 100 nM alectinib or DMSO vehicle control for the indicated times. Conditioned media was collected and assayed for CXCL10/IP-10 with an ELISA kit from Invitrogen and CXCL1, CXCL5/LIX, CCL2 and CCL5/RANTES with ELISA kits from R&D Systems following manufacturer’s instructions. The measured concentration in each sample was normalized to the total cellular protein per dish and the data are presented as pg/μg protein.

### Cell Proliferation Assays

Cell lines were plated at 100 cells per well in 96-well tissue culture plates. 24 hrs later, cells were treated in triplicate with the indicated concentration of alectinib or lorlatinib. Cell number per well by DNA content was determined after 7-10 days of culture using CyQUANT Direct Cell Proliferation Assay (Life Technologies) according to manufacturer’s instructions. Data are presented as percent of control.

### Mice

Wild-type (WT; C57BL/6J; #000664) and green fluorescent protein (GFP)-expressing mice of C57BL/6 strain (C57BL/6-Tg(UBC-GFP)30Scha/J; #0043530) were obtained from Jackson Laboratory (Bar Harbor, ME). *nu/nu* nude mice (Hsd:Athymic Nude-*Foxn1*^*nu*^; #069) were obtained from Envigo (Indianapolis, IN). Experiments were performed in 8-12 week old male and female mice. Animals were bred, housed, and maintained at the University of Colorado Anschutz Medical Campus vivarium. All procedures and manipulations were performed under a protocol approved by the Institutional Animal Care and Use Committee at the University of Colorado Anschutz Medical Campus. Mice were sacrificed using CO_2_ and cervical dislocation as a secondary method.

### Orthotopic Mouse Model of EML4-ALK Lung Cancer

Murine lung cancer cells were injected into the left lobe of the lungs of either C57BL/6J mice or *nu/nu* nude mice. Cells were prepared in a solution of 1.35 mg/mL Matrigel (Corning #354234) diluted in Hank’s Balanced Salt Solution (Corning) for injection. Mice were anesthetized with isoflurane, the left side of the mouse was shaved, and a 1 mm incision was made to visualize the ribs and left lobe of the lung. Using a 30-gauge needle, 5 × 10^5^ cells were injected in 40μL of matrigel cell mix directly into the left lobe of the lung and the incision was closed with staples. Tumors established for 7-10 days and then the mice were submitted to micro-CT (μCT) imaging to obtain pre-treatment tumor volumes. Tumor-bearing mice were randomized into treatment groups (n=10), either 20 mg/kg alectinib or diluent control (H_2_O) by oral gavage 5 days/week until the end of study or termination of treatment. Mice were imaged weekly by μCT imaging to monitor effects of drug treatment on tumor volume. In some experiments, tumor-bearing mice were treated with either H_2_O control or 20 mg/kg alectinib for 4 days. After sacrifice, tumors were measured via digital calipers, and processed for single cell suspensions or formalin fixation.

### Tumor Volume Quantification

μCT imaging was performed by the Small-Animal IGRT Core at the University of Colorado Anschutz Medical Campus in Aurora, CO using the Precision X-Ray X-Rad 225Cx Micro IGRT and SmART Systems (Precision X-Ray, Madison, CT). Tumor volume was quantified from μCT images using ITK-SNAP software (18) (www.itksnap.org).

### Tissue Harvest and Processing

At tissue harvest, lungs were perfused with 5 mL of PBS/heparin (20U/mL, Sigma) and inflated with 4 mL 4% paraformaldehyde (PFA; Electron Microscopy Scientific). The left lung was removed and fixed for 24 hours in 4% PFA, after which they were switched to 70% ethanol. Tissues were processed and embedded into formalin-fixed paraffin-embedded (FFPE) blocks by the Pathology Shared Resource at the University of Colorado Anschutz Medical Campus. Blank slides were cut by the Pathology Shared Resource to be used for multispectral immunofluorescence imaging. To prepare single cell suspensions, mice were sacrificed and the lungs were perfused with 5 mL of PBS/heparin solution. The tumor-bearing left lungs were mechanically dissociated using a razor blade and incubated for 30 minutes at 37°C with 3.2mg/mL Collagenase Type 2 (Worthington, 43C14117B), 0.75 mg/ml Elastase (Worthington, 33S14652), 0.2 mg/ml Soybean Trypsin Inhibitor (Worthington, S9B11099N), and DNAse I 40 μg/ml (Sigma). The resulting single cell suspensions were filtered through 70μm strainers (BD Biosciences), washed with FA3 staining buffer [phosphate-buffered saline (PBS) containing 1% FBS, 2mM EDTA, 10mM HEPES]. Samples underwent a red blood cell lysis step (0.15 mM NH_**4**_Cl, 10 mM KHCO_**3**_, 0.1 mM Na_**2**_EDTA, pH 7.2), were washed, and filtered through a 40 μm strainer (BD Biosciences). Single cell suspensions were then submitted to staining for FACS and flow cytometry. For RNAseq of sorted cancer cell suspensions, lungs from 3-5 mice were pooled together. For flow cytometry, each data point represents a single cell suspension of the left lung of one mouse.

### Fluorescence-Activated Cell Sorting (FACS)

Single cell suspensions were submitted to FACS. Cell sorting was performed at the University of Colorado Cancer Center Flow Cytometry Shared Resource using a MoFlo XDP cell sorter equipped with a 100 micron nozzle (Beckman Coulter). The sorting strategy excluded debris and cell doublets by light scatter and dead cells by DAPI (1 μg/ml). For these studies, lung cancer cells were implanted into GFP+ mice, with cancer cells separated from the host’s GFP-expressing cells by sorting for GFP-negative cells. Immediately after sorting, cells were pelleted and frozen in liquid nitrogen in preparation for RNA extraction. The number of recovered cells ranged from 2.4 × 10^5^ to 15 × 10^5^.

### RNAseq of Cancer Cells Recovered from Tumors

Cancer cells were injected into GFP+ mice and recovered from tumors by FACS as previously described (13,15). Total RNA was isolated from FACS-sorted tumor cells using an RNeasy Plus Mini Kit (QIAgen). The quality and quantity of RNA were analyzed using a bioanalyzer (4150 TapeStation System; Agilent). RNA was submitted to the University of Colorado Cancer Center Genomics Shared Resource for RNAseq library preparation and sequencing using the Illumina HiSEQ2500 (EA1) and NovaSEQ6000 (EA2). Fastq files were analyzed as previously reported (19). Briefly, the Illumina HiSeq Analysis Pipeline was followed. Reads were quality checked using FastQC, aligned with TopHatv2 using the *Mus musculus* mm10 reference genome (UC Santa Cruz), and then the aligned reads were assembled into transcripts using Cufflinks v2.0.2 to estimate their abundance. Data is shown as fragments per kilobase of exon per million fragments mapped (FPKM).

### RNAseq of Murine EML4-ALK Cell Lines

The murine EML4-ALK cell lines cultured in 10 cm dishes were treated for 1-5 days with 0.1% DMSO or 100 nM alectinib. RNA was submitted to the University of Colorado Cancer Center Genomics Shared Resource where libraries were generated and sequenced on the NovaSeq 4000 to generate 2×151 reads. Fastq files were quality checked with FastQC, Illumina adapters trimmed with bbduk, and mapped to the mouse mm10 genome with STAR aligner. Counts were generated by STAR’s internal counter and reads were normalized to counts per million (CPM) using the edgeR R package (20). Heatmaps were generated in Prism 9 (GraphPad Software, San Diego, CA).

### Multispectral Imaging

Methods for multispectral imaging have previously been published and the methods herein are described in brief (21,22). Following the manufacturer’s protocol, FFPE slides were sequentially stained using Opal IHC Multiplex Assay (PerkinElmer, Waltham, MA) by the Human Immune Monitoring Shared Resource at the University of Colorado Anschutz Medical Campus. The Vectra Polaris Imaging System (PerkinElmer) scanned the whole slide. The panel used for the Vectra Polaris were as follows in sequential order: CD11b, CD64, EpCAM. CD11c, B220, CD8, F4/80. Approximately 5 regions of interest (ROIs) were evaluated per tumor. Images were analyzed using inForm Tissue Analysis Software (v2.4.8, Akoya, Menlo Park, CA) to un-mix fluorochromes, remove autofluorescence, segment tissue and cells, and phenotype cells. Data analysis was performed as previously described by our group (13,21). Briefly, data acquired from inForm was analyzed using the R package Akoya Biosciences phenoptrReports. The count_phenotypes function was used to aggregate phenotype counts for each slide. Data are presented as cell count.

### Flow Cytometry

The single cell suspension samples were blocked in anti-mouse CD16/CD32 (clone 93; eBioscience) at 1:200 on a rocker for 15 min at 4°C. Next, fix viability dye (LIVE/DEAD Fixable Aqua Dead Cell Stain Kit; 1:200; Invitrogen) and conjugated antibodies were added (see below) to the single cell suspension. Cells were incubated in the dark at 4°C for 60 minutes. Cells were then resuspended in FA3 buffer and ran on the Gallios Flow Cytometer (Beckman Coulter). For compensation, single-stained beads (VersaComp Antibody Capture Bead Kit; Beckman Coulter) and a single-stained cell-mix of all samples analyzed were used. Flow cytometry was analyzed using Kaluza Analysis Software (v2.0, Beckman Coulter). Compensation was first performed on the single-stained bead controls and then confirmed using the single-stained cell mixture.

#### Antibody Panel

CD11b-FITC (clone M1/70; 1:100; BioLegend), CD64-PE (clone X54-5/7.1; 1:100; BD Biosciences), MHCII-Dazzle (clone M5/111.15.2; 1:250; BioLegend), Ly6C-PerCP/Cy5.5 (clone HK1.4; 1:100; BioLegend), Ly6G-PE/Cy7 (clone 1A8; 1:200; BioLegend), SigF-A647 (clone E50-2440; 1:100; BD Biosciences), CD45-AF700 (clone 30-F11; 1:100; Invitrogen), CD11c-APC/Cy7 (clone HL3; 1:100; BD Biosciences), MHCI-eF450 (clone 28-14-8; 1:100; Invitrogen), CD4-V500 (clone RM4-5; 1:200; BD Biosciences; used only for compensation)

### Mass Cytometry by Time of Flight (CyTOF)

We have previously published this method and statistical analyses for CyTOF (12,23). Briefly, single cell suspensions were treated with benzonase nuclease (#E1014, 1:10,000; Sigma-Aldrich). Cells were stained with cisplatin (Cell-ID Cisplatin, 195Pt, Fluidigm), fixed with 1.6% paraformaldehyde, and snap frozen in 100μL of freezing mix containing PBS, 0.02% Azide, 0.5% BSA, 10% DMSO. After thawing on ice, cells were barcoded with Cell-ID 20-Plex Pd Barcoding Kit (#201060; 102, 104, 105, 106, 108, and 110Pd bar codes; Fluidigm) following the manufacturer’s protocol. Following barcoding, all samples were combined into a single tube for staining. Cells were blocked with anti-mouse CD16/CD32 (24G2; 1:100) for 10 minutes at room temp. Next, cells were stained with primary surface antibodies for 30 minutes at room temp followed by secondary surface antibody staining for 30 minutes at room temp. Cells were then permeabilized overnight following the manufacturere’s protocol for FoxP3 Transcription Factor Fixation/Permeabilization Kit (eBioscience). The following day, the cells were stained with intracellular antibodies for 2 hours at 4°C. Next, cells were fresh fixed with 1.6% PFA (Pierce). Cells were resuspended and incubated with intercalator (191Ir-Intercalator, 193Ir-Intercalator; Cell-ID, Fluidigm) and then run on the Helios Mass Cytometer (Fluidigm) in the University of Colorado Cancer Center Flow Cytometry Shared Resource. All antibodies were stained at a dilution of 1:100. *Primary surface antibodies used*: 89Y-CD45 (Clone 30-F11; Fluidigm), 141Pr-Gr1 (Ly6C/Ly6G; Clone RB6-8C5; Fluidigm); 142Nd-CD11c (Clone N418; Fluidigm), 143Nd-GITR (Clone DTA1; Fluidigm), 144Nd-MHCI (Clone 28-14-8; Fluidigm), SiglecF-PE (Clone E50-2440; BF), 146Nd-CD8a (Clone 53-6.7; Fluidigm), 148Nd-CD11b (Clone M1/70; Fluidigm), 149Sm-CD19 (Clone 6D5; Fluidigm), 150Nd-CD25 (Clone3C7; Fluidigm), 151Eu-CD64 (Clone X54-5/7.1; Fluidigm), 152Sm-CD3e (Clone 145-2C11; Fluidigm), 153Eu-PD-L1 (Clone 10F.9G2; Fluidigm), 154Sm-CTLA4 (Clone UC10-4B9; Fluidigm), 156Gd-CD90.2 (Clone 30-H12; Fluidigm), 159Tb-PD-1 (Clone RMP1-30; Fluidigm), 162Dy-Tim3 (Clone RMT3-23; Fluidigm), CXCR3-APC (Clone CXCR3-173; BioLegend), 167Er-NKp46 (Clone 9A1.4; Fluidigm), 169Tm-Ly-6A/E (Sca-1; Clone D7; Fluidigm), CD103-Biotin (Clone 2E7; BioLegend), 171Yb-CD44 (Clone IM7; Fluidigm), 172Yb-CD4 (Clone RM4-5; Fluidigm), 173Yb-CD117 (c-kit; Clone 2B8; Fluidigm), 174Yb-Lag3 (Clone M5/114.15.2; Fluidigm), 175Lu-CD127 (Clone A7R34; Fluidigm), 176Yb-ICOS (Clone 7E.17G9; Fluidigm), 209Bi-MHCII (I-A/I-E; Clone M5/114/15/2; Fluidigm). *Secondary surface antibodies used:* 145Nd-PE (Clone PE-001; Fluidigm), 163Dy-APC (Clone APC003; Fluidigm), 170Er-Biotin (Clone 1D4-C5; Fluidigm). *Intracellular antibodies used:* 147Nd-p-H2AX [Ser139] (Clone JBW301; Fluidigm), 155Gd-IRF4 (Clone 3E4; Fluidigm), 158Gd-FoxP3 (Clone FJK-16s; Fluidigm), 161Dy-iNOS (Clone 4B10; Fluidigm), 164Dy-IkBa (Clone L35A5; Fluidigm), 165Ho-Beta-catenin (Clone D13A1; Fluidigm), 166Er-Arginase1 (Clone 6D5; Fluidigm), 168Er-Ki-67 (Clone Ki-67; Fluidigm). Normalization Beads were used (140Ce, 151Eu, 153Eu, 165Ho, 175Lu). Each sample represents a tumor-bearing left lung from one mouse.

#### CyTOF Analysis

The raw FCS files acquired on the Helios were first normalized using the Matlab Bead Normalizer downloaded from the Nolan laboratory GitHub page (NormalizerR2013a(8.1)_64bitWindows; https://github.com/nolanlab). The normalized files were next debarcoded using SingleCellDebarcoderR2013a(8.1)_64bitWindows that was downloaded from the Nolan laboratory GitHub page (https://github.com/nolanlab). Normalized/debarcoded data were gated for live singlets events (191Ir+/193Ir+/195Pt-) using traditional Boolean gating FlowJo software (v10.8.1; Ashland, OR), and FCS files were downloaded for biaxial gating and analyses using Kaluza Analysis Software.

### Graphical Analysis and Statistics

Graphing and statistical analysis was performed using GraphPad Prism version 9.2.0 (San Diego, CA) or R version 4.0.3, R studio (for multispectral imaging analyses). To determine significance between groups, student’s t-tests or one-way ANOVA tests were performed. Data are presented as mean ± SEM. Significant differences are indicated by *p<0.05, **p<0.01, ***p<0.001, or ****p<0.0001.

### Data Availability

The RNAseq data generated in this study are being deposited in the GEO Datasets.

## Results

### Therapeutic response of primary EML4-ALK tumors to ALK-targeting TKIs

Maddalo et al (17) demonstrated that intratracheal delivery of an adenovirus encoding Cas9 and gRNAs targeting murine *Eml4* and *Alk* yielded efficient generation of EML4-ALK positive murine lung tumors. We observed that instillation of Adeno-Cas9-gRNA virus induces multiple tumor foci in C57BL/6 mice (**Supplementary Figure S1**). Although limited by the sensitivity of μCT as a detection method, the vast majority of lesions underwent rapid and significant shrinkage following TKI treatment as initially noted (17). Moreover, TKI treatment for 4 weeks and then termination of therapy resulted in prompt regrowth of the lesions within 4 weeks, indicating the presence of residual disease. Thus, this murine model of EML4-ALK-driven lung cancer replicates a key feature of the human disease. For further investigations of the molecular mechanisms mediating variation in the depth and duration of response, we developed a panel of transplantable murine EML4-ALK cancer cell lines derived from this model.

### *Murine EML4-ALK positive lung cancer cell lines exhibit equivalent* in vitro *sensitivity to TKIs, but distinct therapeutic responses in an orthotopic model*

A murine ALK+ cell line referred to herein as EA2 is a generous gift of Dr. Andrea Ventura (17). We also developed two additional EML4-ALK-driven cell lines, EA1 (15) and EA3 within our institution. All cell lines were isolated from C57BL/6 mice and express the mRNA encoding the EML4-ALK variant 1 fusion gene compared to KRAS-mutant murine LLC lung cancer cells (**Supplementary Figure S2A and S2B**). EA2 and EA3 cell lines were derived from TP53 null mice while EA1 was derived from a TP53 wild-type mouse. The immunoblot in **Supplementary Figure S2C** demonstrates TP53 protein expression in EA1, but not EA2 or EA3 cells.

The three cell lines exhibited equivalent *in vitro* growth sensitivity to the ALK TKIs, alectinib and lorlatinib (**Figure 1A**). In addition, the findings in **Figure 1B** show equivalent reduction of mRNA levels of the MAPK pathway-regulated genes, ETV4, ETV5, FOSL1 and DUSP6 (24,25) by alectinib. Thus, these data demonstrate equal alectinib sensitivity of EA1, EA2 and EA3 cells using functional *in vitro* assays.

**Figure 1.**
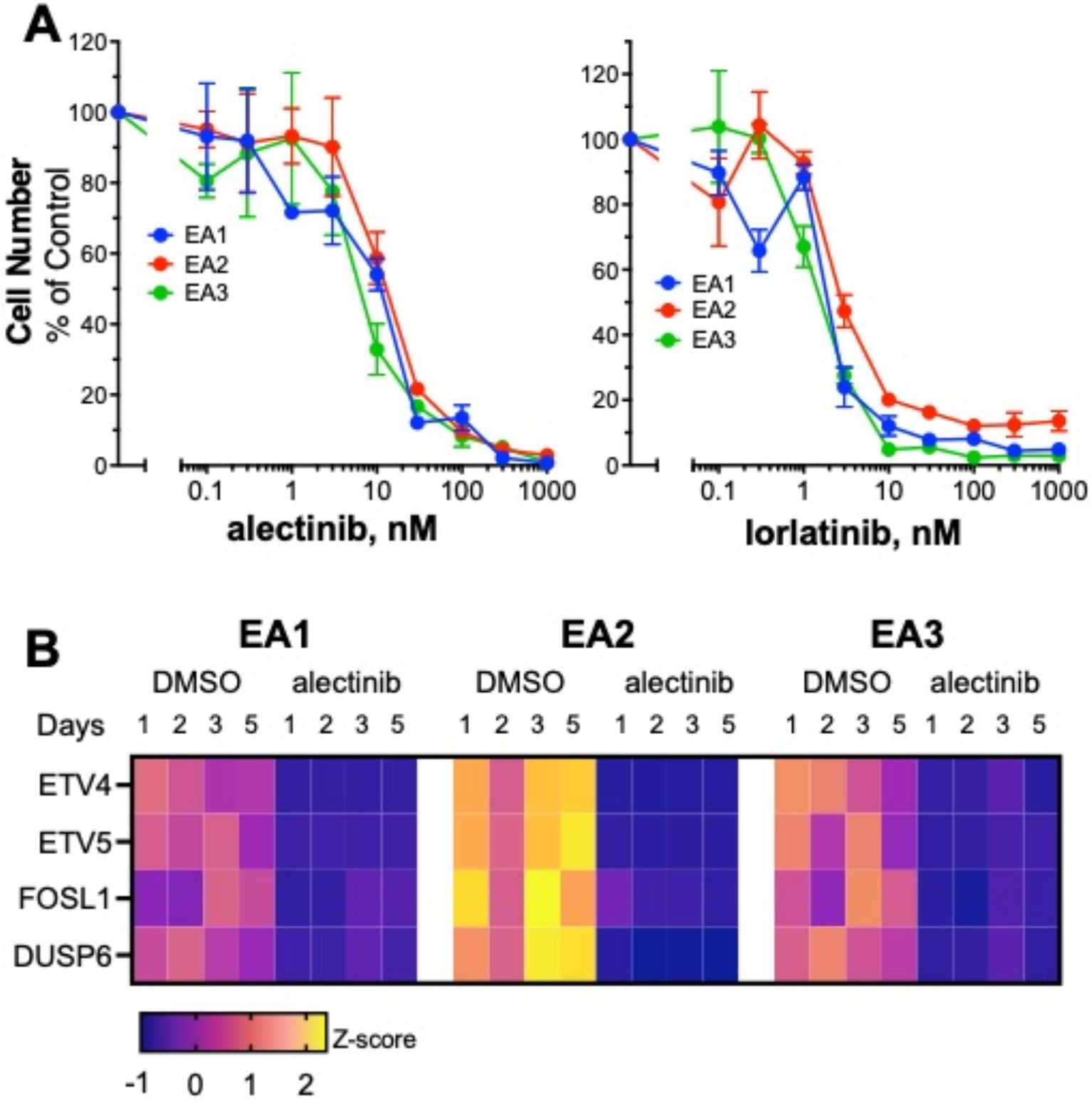
Murine EML4-ALK cell lines exhibit equal *in vitro* sensitivity to TKIs. **A**, EA1, EA2 and EA3 cells were seeded at 100 cells/well in 96-well plates and cultured in growth medium containing the indicated concentrations of the TKIs, alectinib or lorlatinib. After 7 days of incubation, cell number was determined using the CyQUANT assay (see Materials and Methods). Data (mean and SEM of triplicate determinations) are presented as percent of wells treated with medium containing 0.1% DMSO. The IC_50_ values for alectinib (7.6, 12.4 and 6.7 nM for EA1, EA2 and EA3, respectively) and lorlatinib (1.7, 3.8 and 1.7 nM for EA1, EA2 and EA3, respectively) were calculated with Prism 9. **B**, RNAseq data from the murine EML4-ALK cell lines treated with and without alectinib (see Materials and Methods) were queried for known MAPK target genes, ETV4, ETV5, FOSL1 and DUSP6. The mRNA expression values (CPMs) were converted to Z-scores and presented as a heatmap.

The three EML4-ALK cell lines were used to establish orthotopic tumors in the left lung lobe of syngeneic C57BL/6 mice (see Materials and Methods and (12,14,26,27)). Following tumor establishment for ∼10 days, the initial tumor volume was measured by micro CT (μCT), and the mice were treated daily with alectinib (20 mg/kg) or diluent by oral gavage. Tumor volume was measured weekly by μCT. Marked tumor shrinkage was induced by alectinib in all three models **(Figure 2)**. Notably, residual disease was detectable after 3 weeks of continuous treatment in EA1 and EA3 tumors while the EA2 tumors yielded a significantly greater alectinib-mediated tumor shrinkage such that residual disease could not be detected by μCT. Following alectinib treatment termination, EA1 and EA3-derived tumors rapidly regrew within 2 to 3 weeks to volumes equal to the control diluent-treated mice (**Figure 2**), confirming the presence of residual disease. By contrast, no regrowth was observed in EA2 tumors 5 weeks after stopping treatment. The data reveal that EA2 tumors exhibit a complete response to alectinib whereas EA1 and EA3 tumors exhibit partial responses.

**Figure 2.**
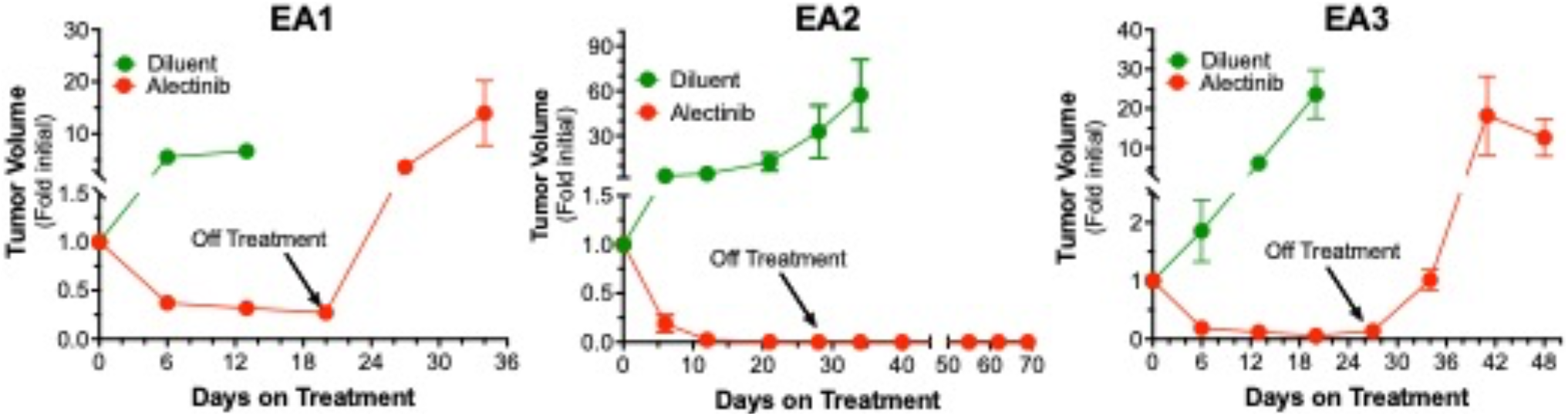
Alectinib responsiveness of orthotopically-propagated EML4-ALK tumors in syngeneic mice. 500,000 cells of each cell line were implanted into the left lungs of C57BL/6 mice. The orthotopic tumors established for ∼10 days, the mice were submitted to μCT imaging to determine pre-treatment tumor volumes and following randomization, treated daily with 20 mg/kg alectinib or diluent by oral gavage. Mice were imaged by μCT weekly over the course of the experiments and tumor volume is presented as the fold change from the initial pre-treatment measurement. The initial tumor volumes (mean ± SEM) for the diluent and alectinib-treated groups were 8.5±7.9 mm^3^ and 11.3±9.1 mm^3^, 10.9±8.5 mm^3^ and 11.6±8.8 mm^3^ and 2.6±2.5 mm^3^ and 1.9±1.6 mm^3^ for EA1, EA2 and EA3, respectively. Where indicated by arrows, alectinib treatment was terminated and assessment of tumor volume by μCT was continued.

### Alectinib induces an IFNγ-like transcriptional program and varied expression of distinct chemokines

Our recent studies in EGFR mutant lung cancer cell lines and murine head and neck cancer cells demonstrated a TKI-stimulated IFNγ transcriptional response accompanied by increased chemokine expression (7,28). The three murine EML4-ALK cell lines were treated *in vitro* with alectinib or DMSO and RNAseq was performed (see Materials and Methods). The resulting datasets were submitted to gene-set enrichment analysis (GSEA) using the Hallmark Pathways and the normalized enrichment scores (NES) are presented as a heatmap in **Supplementary Figure S3**. The data show that multiple inflammation-related Hallmark pathways are enriched in alectinib-treated cells including IFNα, IFNγ, allograft rejection and inflammatory response. In addition, Hallmark pathways associated with cell proliferation (MYC targets V1 and V2, G2M checkpoint, E2F targets) are markedly de-enriched in the alectinib-treated cells and provides further support for equivalent growth inhibition by alectinib in the three cell lines.

To interrogate diversity of chemokine expression amongst the EML4-ALK lines when propagated as orthotopic tumors, the cancer cells from untreated EA1 and EA2 tumors were purified from transgenic GFP-expressing mice and submitted to RNAseq analysis (see Materials and Methods). As shown in **Figure 3**, mRNA levels of CXCL9 and CXCL10, two chemokines with T cell recruitment functions, were significantly higher in EA2 tumors compared to EA1. CCL2 and CCL7 expression was also higher in untreated EA2 relative to EA1 tumors. By contrast, levels of CXCL1, CXCL2, CXCL12 and TGF-β2, chemokines and cytokines associated with recruiting immunosuppressive populations (29) were higher in EA1 tumors relative to EA2 tumors.

**Figure 3.**
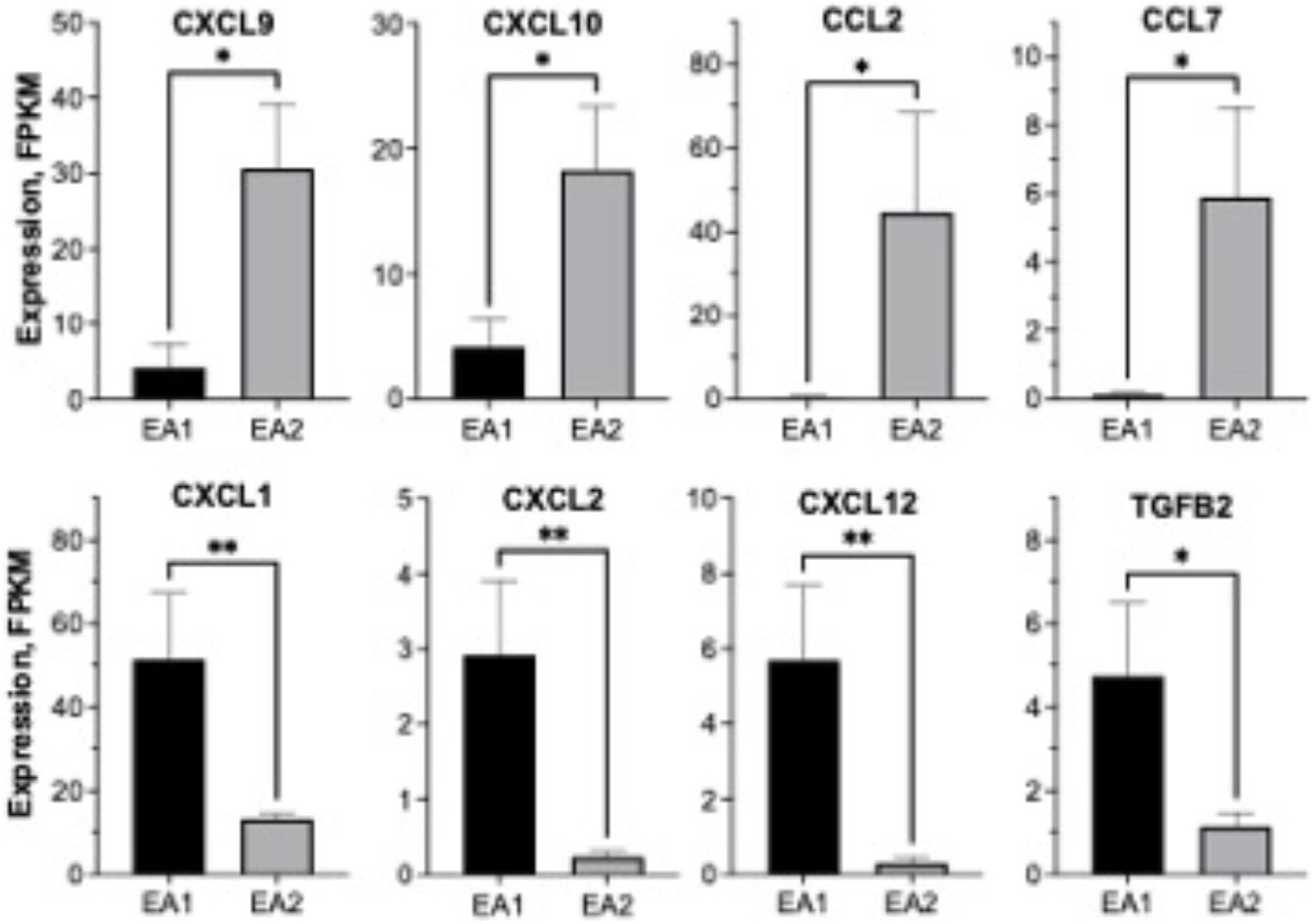
Distinct baseline chemokine expression patterns in EA1 and EA2 orthotopic tumors. EA1 and EA2 tumors were established in C57BL/6 mice in which GFP was ubiquitously-expressed in all cell types (see Materials and Methods). After ∼3 weeks, the left lung lobes from the tumor bearing mice (n=5 for each cell line) were dissociated into single-cell suspensions and submitted to flow sorting to collect the GFP-negative tumor cells. RNA was purified from the sorted cells and submitted to RNAseq. The data show the mRNA expression in FPKM (mean and SEM) where * and ** indicate p-values less than 0.05 and 0.01 respectively.

Specific chemokines and cytokines are central to the genes comprising the inflammation-related Hallmark pathways. Using the RNAseq data as a guide, multiple chemokine genes were selected and the effect of alectinib treatment on their secretion was determined by ELISA. The findings in **Figure 4** reveal alectinib-stimulated secretion of multiple chemokines with diverse activities as anti- and pro-tumorigenic factors. CXCL10 and CCL5 were markedly induced in response to alectinib in EA2 and EA3 cells, but weakly in EA1 cells. By contrast, alectinib-induced secretion of CXCL1, CXCL5 and CCL2 with roles in the recruitment of lymphocytes of myeloid lineages were similar among the three cell lines. In general, the level of chemokine expression measured *in vitro* with untreated cell lines was very low, indicating that signals from the murine hosts are distinctly integrated by EA1 and EA2 cells to achieve the distinct levels observed *in vivo*.

**Figure 4.**
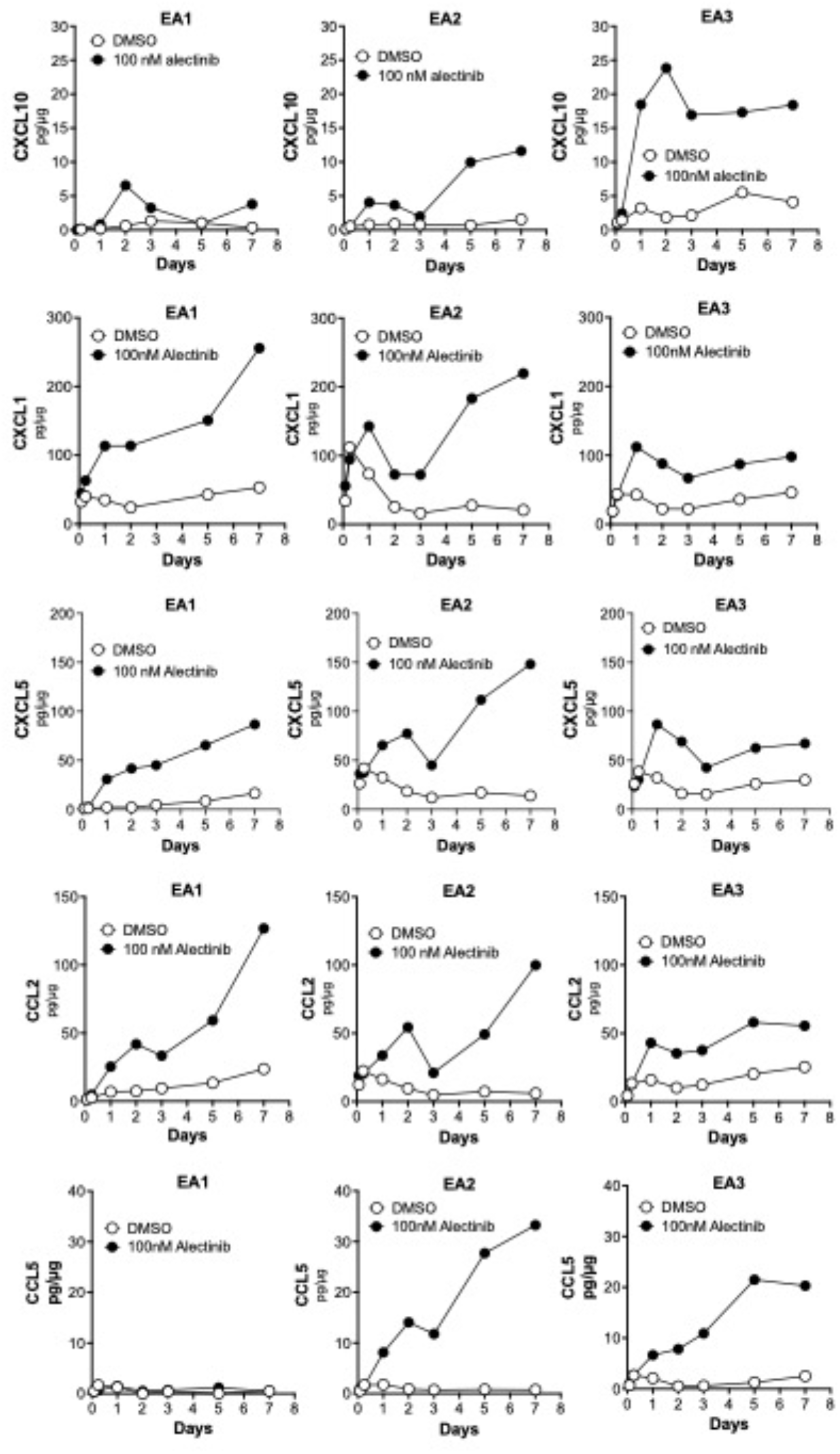
Chemokine induction with alectinib treatment in murine EML4-ALK cell lines. EA1, EA2 and EA3 cells were treated *in vitro* with 0.1% DMSO or 100 nM alectinib for 2 hours to 7 days. The culture medium was collected and submitted to ELISA for the indicated chemokines. The data are presented as pg of analyte per μg cell protein and are representative of 2 independent experiments.

### Analysis of the baseline and TKI-regulated immune microenvironment in murine EML4-ALK tumors

The findings in **Figures 3** and **4** and **Suppl. Fig. S3** support an emerging view that oncogene-targeted drugs regulate cancer cell-derived signaling via cytokines and chemokines to the host immune microenvironment (9-11,30,31). Moreover, a predicted consequence of intra-tumoral variation in baseline and TKI-induced chemokine expression is distinct immune cell compositions of the respective tumor microenvironments. To analyze the immune cell populations in control and alectinib-treated tumors, a combination of multispectral imaging (Polaris Vectra), CyTOF and standard flow cytometry was deployed with the EA1 model which exhibits a partial response to alectinib and the EA2 model that exhibits a complete response (**Figure 2**).

#### Baseline immune cell composition in EA1 and EA2 tumors

Left lungs from mice bearing untreated EA1 and EA2 tumors were dissociated into a single cell suspension and submitted to CyTOF analysis with a panel of 36 antibodies labeled with heavy metal isotopes. The resulting datasets were submitted to gating strategies that revealed increased CD45+ cells in EA2 tumor bearing lungs. Differences in the frequency of CD4+ and CD8+ T cells or B cells in EA1 and EA2 tumor-bearing lungs were not observed (**Supplementary Figure S4**). However, increased baseline content of alveolar macrophages (MacA: defined as CD11c+/SigF+) and recruited monocyte/macrophage populations (MacB: defined as CD11b+/Ly6G-/SigF-) (19) were observed in EA2 lungs and tumors relative to EA1 tumors, with a trend towards increased baseline content of neutrophils (p=0.078) in EA1 tumors. The findings from CyTOF analysis demonstrate unique cellular distribution between EA1 and EA2 tumors, and are consistent with observed differences in baseline chemokine expression between these tumors (**Figure 3**).

#### Alectinib-induced Immune cell changes in control and alectinib-treated EA1 and EA2 tumors

EA1 and EA2 cells were implanted into mice and allowed to establish for 3 weeks and then treated for 4 days with control or 20 mg/kg alectinib. The left lungs of C57BL/6 mice bearing established EA1 and EA2 tumors were harvested, formalin-fixed and submitted to Polaris Vectra imaging with a panel of antibodies targeting immune cells (see Materials and Methods). We focused on CD8+ T cells, as defined as CD3+/CD8+ positivity, and the CD3+/CD8-phenotype identifying CD4+ T cells. Also, B cells were assessed by positivity for B220 staining. In untreated tumors, a significant increase in the number of tumor-infiltrated CD8+ cells was observed in EA2 tumors compared to EA1, with comparable numbers of CD3+/CD8-cells (**Figure 5A**). Following 4 days of alectinib treatment, EA2 tumors exhibited a trend towards further increase in CD8+ T cells, although not statistically significant, while EA1 tumors showed no change in CD8+ or CD4+ T cell content with treatment.

**Figure 5.**
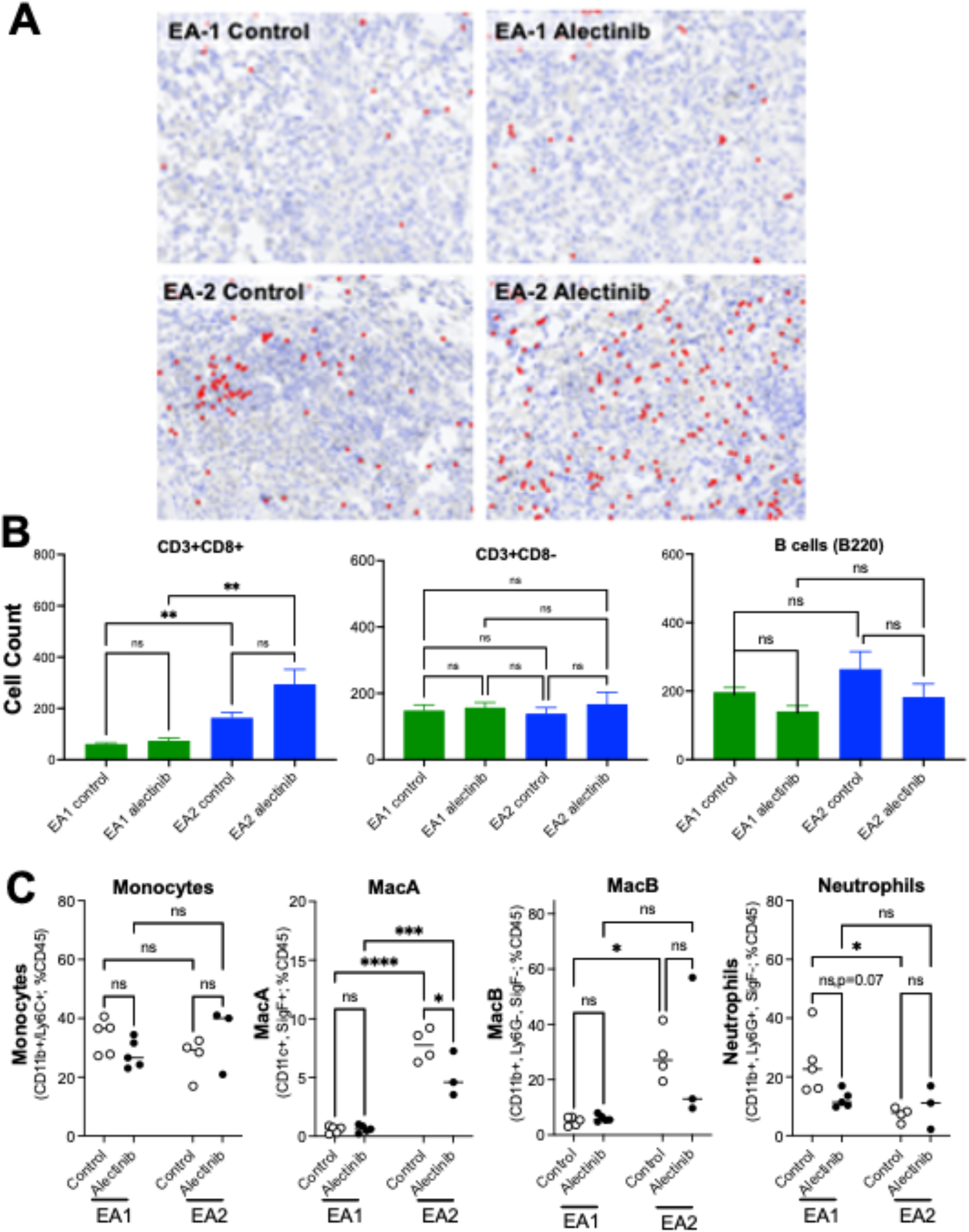
Adaptive and innate immune cells in control and alectinib-treated EA1 and EA2 tumors. EA1 or EA2 cells were implanted in the left lung of C57BL/6 mice. Tumors were established for 3 weeks and then mice were treated for 4 days with either control or 20 mg/kg alectinib. **A**, Tissue was harvested, processed, and FFPE slides were submitted to multispectral imaging using Vectra Polaris. Three to five regions of interest were quantified for each tumor. Data acquisition, segmentation, and cell phenotyping was performed using inForm software. The R package Akoya Biosciences phenoptrReports was used to count CD3+/CD8+ and CD3+/CD8-cells (presumed to be CD4+ T cells) as well as B220-positive B cells. Differences between groups was analyzed by the Kruskal-Wallis test with correction for multiple comparisons. **B**, After 4 days of treatment, mice were sacrificed and a single-cell suspension was made from the left tumor-bearing lung. The single cell suspension was stained and submitted to flow cytometry analysis for innate immune cells using the previously described gating strategy (19). Differences between groups was analyzed with ordinary one-way ANOVA and correction for multiple comparisons. Data are presented as the mean ± SEM were *, ***, and **** indicate p-values less than 0.05, 0.001, and 0.0001, respectively.

To identify potential changes in myeloid cell types, we submitted dissociated tumor-bearing lungs to flow cytometric analysis using a previously reported gating strategy (19). No differences in monocyte (defined as CD11b+/Ly6C+) content was observed between EA1 and EA2 tumor-bearing lungs harvested from control or alectinib-treated mice (**Figure 5B**). Control EA2 tumors exhibited increased content of MacA and MacB cells compared to EA1 tumors, which did not significantly change with a 4-day alectinib treatment (**Figure 5B**). By contrast, control EA1 tumors had elevated neutrophil (defined as CD11b+/Ly6G+/SigF-) content (consistent with **Suppl. Fig. S4**) compared to EA2 tumors and alectinib led to a decrease that approached statistical significance in the neutrophil population in EA1 tumors (**Figure 5B**).

### Adaptive immunity is required for durable alectinib responses

In our previous study, pan-ERBB TKI response in orthotopic murine head and neck tumors was diminished in immune-deficient hosts (28). To test the role of the adaptive immune system in the alectinib therapeutic response shown in **Figure 2**, the murine EML4-ALK cells were inoculated into the left lungs of *Nu/Nu* mice lacking functional T and B cells. In all three EML4-ALK cell line models, significant tumor shrinkage was elicited by alectinib, but was followed by prompt tumor progression within three to four weeks of initiating treatment, despite continuous TKI therapy (**Figure 6**). These data indicate that functional adaptive immune cells are critical for the sustained anti-cancer activity of alectinib observed in C57BL/6 mice (**Figure 2**).

**Figure 6.**
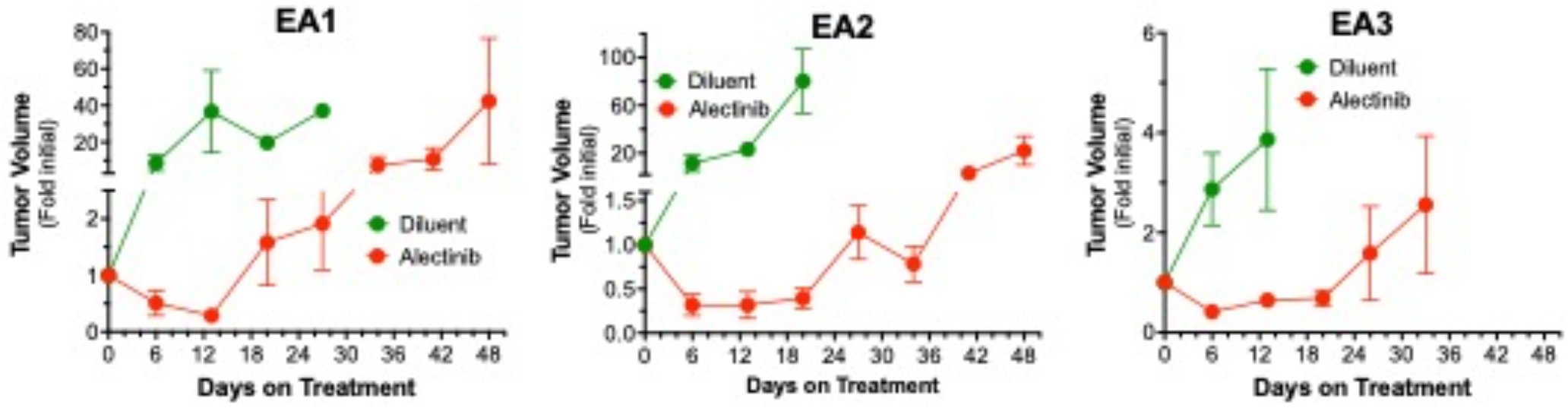
Alectinib responsiveness of orthotopically-propagated EML4-ALK tumors in immune-deficient *nu/nu* mice. 500,000 cells of each cell line were implanted into the left lungs of *nu/nu* mice. The orthotopic tumors established for ∼10 days, the mice were submitted to μCT imaging to determine pre-treatment tumor volumes and following randomization, treated daily with 20 mg/kg alectinib or diluent by oral gavage over the entire course of the experiment. Mice were imaged by μCT weekly over the course of the experiments and tumor volume is presented as the fold change from the initial pre-treatment measurement. The initial tumor volumes (mean ± SEM) for the diluent and alectinib-treated groups were 3.8±2.6 mm^3^ and 4.2±3.7 mm^3^, 8.7±10.5 mm^3^ and 9.4±11.2 mm^3^ and 18.9±11.6 mm^3^ and 20.5±13.5 mm^3^ for EA1, EA2 and EA3, respectively.

## Discussion

A major finding of this study is that durable responses to alectinib in the murine EML4-ALK tumor models require a functional adaptive immune system. The present study builds on the findings in our recent report showing that the degree of induction of an IFNγ transcriptional program measured in on-treatment biopsies obtained after ∼2 weeks of treatment with EGFR-specific TKIs positively associated with the duration of the therapeutic response (7). In the Gurule et al study, a transcriptional signature for T cells was increased in the on-treatment biopsies from patients exhibiting longer treatment duration, suggesting a role for T cells in the duration of response. Similarly, we have shown that pan-ERBB inhibitor-induced tumor shrinkage of a murine EGFR-dependent head and neck cancer model was markedly reduced in *nu/nu* mice compared to syngeneic BALB/c mice (28). In this regard, there is a growing body of evidence that oncogene-targeted agents induce cancer cell-derived signals instructing functional interaction of the adaptive and innate immune systems (8,10,11,30,32). Canon et al demonstrated that *in vivo* treatment of a murine KRAS-G12C-driven tumor model with a KRAS-G12C inhibitor resulted in a pro-inflammatory TME and the duration of the therapeutic response was greatly truncated in immune-deficient hosts (33). Using scRNA analysis of human biopsies, Maynard et al showed a transient immunostimulatory effect in EGFR-mutant patients after initial TKI therapy, followed by establishment of an immunosuppressive environment upon progression (22) While we were not able to assess potential variation in duration of treatment among the three murine EML4-ALK models as it would likely require many months or more of daily oral delivery of alectinib to tumor-bearing mice, we postulate that variation in the engagement of adaptive immunity with individual tumors may account for the range of therapy durations observed in oncogene-defined subsets of lung cancer patients.

The results in **Figures 3** and **5** indicate that TME-derived signals and TKI-induced signaling represent distinct inputs into the integrated chemokine expression program that presumably drives immune cell infiltration and content. Notably, EA2 tumors exhibiting a complete therapeutic response to alectinib present with increased expression of the T cell-attracting chemokines, CXCL9 and CXCL10, and CD8+ T cell content at baseline relative to EA1 tumors which exhibit a partial response with viable residual disease. Moreover, EA2 cells exhibit a greater *in vitro* induction of CXCL10 upon alectinib treatment compared to EA1 cells (**Figure 4**) and show a trend for further increases in CD8+ T cell content following alectinib treatment *in vivo* (**Figure 5**). By contrast, CXCL1, CXCL2, CCL2 and TGFB2 with defined pro-tumorigenic roles are more highly expressed by EA1 tumor cells at baseline and may antagonize anti-tumorigenic effects of CD8+ T cells that do infiltrate this model. We propose that differential expression of anti- and pro-tumorigenic chemokines by cancer cells is mediated through inherent differences in the cancer cells, as well as their response to signals emanating from the TME. While these mechanisms are not well understood, difference in signaling pathways, epigenetic regulation of specific chemokines and differences in the repertoire of cell surface receptors on different cancer cells are likely to dictate the expression pattern of cytokines and chemokines. In a recent report, Tang et al (34) demonstrated that treatment of KRAS and EGFR-driven murine lung cancer models with a SHP2 inhibitor increased expression of chemokines that promoted infiltration of T and B cells, but also granulocyte myeloid derived suppressor cells (gMDSC’s) via CXCR1/2 ligands. Combined treatment with SHP2 inhibitor and CXCR1/2 antagonists yielded greater therapeutic benefit compared to SHP2 inhibitor alone. Thus, the murine EML4-ALK cells lines and the orthotopic model are predicted to provide deep insight into the complex regulation of the variable immune microenvironment that is established in individual tumors and serve as a robust platform to test various strategies in experimental therapeutics.

The orthotopic implantation model and the panel of murine EML4-ALK cell lines replicate a key feature of the human disease, variable residual disease with continuous alectinib treatment. EA1 and EA3-derived tumors maintain clear residual tumor that can be detected by μCT and drives rapid tumor progression upon termination of therapy. By contrast, EA2 tumors shrink such that no tumor can be visualized with μCT and tumors fail to re-grow upon terminating therapy. Inspection of the alectinib response in mice bearing multiple primary EML4-ALK tumors induced by the Cas9 and gRNA-encoding adenovirus demonstrates that most, but not all of the lesions re-grow following termination of alectinib therapy (**Suppl. Fig. S1**) and indicates that the EA2 phenotype is not simply a result of *in vitro* culture. Whether this presentation of therapeutic variation is the result of heterogeneous contribution from host immunity or is mediated by cancer cell autonomous mechanisms, or perhaps both processes, remains a subject for further experimentation. We presume that the rapid outgrowth of all three cell lines when propagated in immunodeficient *nu/nu* mice in the setting of continuous alectinib treatment is mediated by rapid activation of bypass signaling pathways. The observation that outgrowth does not occur in immunocompetent hosts when on TKI treatment suggests that the adaptive immune system actively performs surveillance of the residual tumor cells to block their outgrowth, potentially monitoring additional TKI-induced changes to tumor cells (e.g. chemokines, antigen presentation machinery). Perhaps the initial reduction of cancer cells alters the balance between T cells and cancer cells as proposed in the immunoediting model of cancer (35) and prevents the ability of cancer cells undergoing bypass signaling to grow. Of note, variant calling algorithms indicate that EA2 cells may bear a 3-fold higher mutation burden relative to EA1 and EA3 (not shown). Alternatively, it is possible that T cells are capable of negatively regulating bypass signaling mechanisms directly. Additional studies are required to dissect these distinct possibilities.

Finally, these data suggest that ALK+ tumors with greater T cell content prior to initiation of therapy may be predicted to exhibit a stronger and potentially longer-lasting response to targeted agents. In support, a clinical study reported improved overall survival of EGFR and ALK patients whose tumors presented with high CD8+ T cells prior to initiating TKI therapy (36). Thus, strategies to increase T cell infiltration and/or functionality in combination with precision oncology agents may represent an approach to improve the depth and duration of response in oncogene-defined subsets of lung cancer patients.

## Authors’ Disclosures

EL Schenk reports speaker fees from the American Lung Association, American Society of Clinical Oncology, OncLive, Physicians Education Resource, Takeda, and Roche/Genentech. EL Schenk reports consultant fees from Actinium, Bionest Partners, ExpertConnect, FCB Health, Guidepoint Network, the KOL Connection Ltd, and the ROS1ders. EL Schenk is on the Scientific Advisory Board for Regeneron and Janssen. EL Schenk reports research funding from Takeda. T Patil is on the advisory boards for Astrazeneca, Pfizer, EMD Soreno, Janssen, Mirati Therapeutics, Sanofi, and Takeda. T Patil has studies that are sponsored by EMD Soreno and Janssen. T Patil has no research funding to disclose. No disclosures were reported by the other authors.

## Authors’ Contributions

Conception and design: E. Kleczko, E. Clambey, M. Weiser-Evans, E. Schenk, R. Nemenoff, L. Heasley

Development of methodology: E. Kleczko, T. Hinz, A. Le, A. Johnson, L. Heasley, R. Nemenoff

Acquisition of data: E. Kleczko, J. Kwak, T. Nguyen, N. Gurule, T. Hinz, A. Navarro, A. Johnson, D. Polhac, E. Schenk, L. Heasley

Analysis and interpretation of data: E. Kleczko, T. Hinz, T. Nguyen, N. Gurule, A. Navarro, A. Johnson, D. Polhac, E. Schenk, L. Heasley, R. Nemenoff

Writing, review, and/or revision of the manuscript: E. Kleczko, L. Heasley, R. Nemenoff, T. Patil, E. Schenk

## Grant Support

This research was supported by the Department of Defense Lung Cancer Research Program award W81XWH1910220 (LEH and RAN), and the University of Colorado Cancer Center Core Grant P30 CA046934.

## Acknowledgments

The authors acknowledge the Genomics and Pathology shared resources within the University of Colorado Cancer and the Human Immune Monitoring Shared Resource within the University of Colorado School of Medicine.

## Notes

### Competing Interest Statement

The authors have declared no competing interest.

## References

1. Camidge DR, Bang YJ, Kwak EL, Iafrate AJ, Varella-Garcia M, Fox SB, et al. Activity and safety of crizotinib in patients with ALK-positive non-small-cell lung cancer: updated results from a phase 1 study. The lancet oncology 2012;13(10):1011–9 doi 10.1016/S1470-2045(12)70344-3.

2. Shaw AT, Gandhi L, Gadgeel S, Riely GJ, Cetnar J, West H, et al. Alectinib in ALK-positive, crizotinib-resistant, non-small-cell lung cancer: a single-group, multicentre, phase 2 trial. The lancet oncology 2016;17(2):234–42 doi 10.1016/S1470-2045(15)00488-X.

3. Bivona TG, Doebele RC. A framework for understanding and targeting residual disease in oncogene-driven solid cancers. Nat Med 2016;22(5):472–8 doi 10.1038/nm.4091.

4. Sharma SV, Lee DY, Li B, Quinlan MP, Takahashi F, Maheswaran S, et al. A chromatin-mediated reversible drug-tolerant state in cancer cell subpopulations. Cell 2010;141(1):69–80 doi 10.1016/j.cell.2010.02.027.

5. Wu W, Haderk F, Bivona TG. Non-Canonical Thinking for Targeting ALK-Fusion Onco-Proteins in Lung Cancer. Cancers (Basel) 2017;9(12) doi 10.3390/cancers9120164.

6. McCoach CE, Blumenthal GM, Zhang L, Myers A, Tang S, Sridhara R, et al. Exploratory analysis of the association of depth of response and survival in patients with metastatic non-small-cell lung cancer treated with a targeted therapy or immunotherapy. Annals of oncology : official journal of the European Society for Medical Oncology / ESMO 2017;28(11):2707–14 doi 10.1093/annonc/mdx414.

7. Gurule NJ, McCoach CE, Hinz TK, Merrick DT, Van Bokhoven A, Kim J, et al. A tyrosine kinase inhibitor-induced interferon response positively associates with clinical response in EGFR-mutant lung cancer. NPJ Precis Oncol 2021;5(1):41 doi 10.1038/s41698-021-00181-4.

8. Kumagai S, Koyama S, Nishikawa H. Antitumour immunity regulated by aberrant ERBB family signalling. Nat Rev Cancer 2021;21(3):181–97 doi 10.1038/s41568-020-00322-0.

9. Petrazzuolo A, Perez-Lanzon M, Martins I, Liu P, Kepp O, Minard-Colin V, et al. Pharmacological inhibitors of anaplastic lymphoma kinase (ALK) induce immunogenic cell death through on-target effects. Cell Death Dis 2021;12(8):713 doi 10.1038/s41419-021-03997-x.

10. Sisler DJ, Heasley LE. Therapeutic opportunity in innate immune response induction by oncogene-targeted drugs. Future Med Chem 2019;11(10):1083–6 doi 10.4155/fmc-2018-0292.

11. Gurule NJ, Heasley LE. Linking tyrosine kinase inhibitor-mediated inflammation with normal epithelial cell homeostasis and tumor therapeutic responses. Cancer Drug Resist 2018;1:118–25 doi 10.20517/cdr.2018.12.

12. Bullock BL, Kimball AK, Poczobutt JM, Neuwelt AJ, Li HY, Johnson AM, et al. Tumor-intrinsic response to IFNgamma shapes the tumor microenvironment and anti-PD-1 response in NSCLC. Life Sci Alliance 2019;2(3) doi 10.26508/lsa.201900328.

13. Johnson AM, Bullock BL, Neuwelt AJ, Poczobutt JM, Kaspar RE, Li HY, et al. Cancer Cell-Intrinsic Expression of MHC Class II Regulates the Immune Microenvironment and Response to Anti-PD-1 Therapy in Lung Adenocarcinoma. J Immunol 2020;204(8):2295–307 doi 10.4049/jimmunol.1900778.

14. Li HY, McSharry M, Bullock B, Nguyen TT, Kwak J, Poczobutt JM, et al. The Tumor Microenvironment Regulates Sensitivity of Murine Lung Tumors to PD-1/PD-L1 Antibody Blockade. Cancer Immunol Res 2017;5(9):767–77 doi 10.1158/2326-6066.CIR-16-0365.

15. Kwak JW, Laskowski J, Li HY, McSharry MV, Sippel TR, Bullock BL, et al. Complement Activation via a C3a Receptor Pathway Alters CD4(+) T Lymphocytes and Mediates Lung Cancer Progression. Cancer Res 2018;78(1):143–56 doi 10.1158/0008-5472.CAN-17-0240.

16. Nolan K, Verzosa G, Cleaver T, Tippimanchai D, DePledge LN, Wang XJ, et al. Development of syngeneic murine cell lines for use in immunocompetent orthotopic lung cancer models. Cancer Cell Int 2020;20:417 doi 10.1186/s12935-020-01503-5.

17. Maddalo D, Manchado E, Concepcion CP, Bonetti C, Vidigal JA, Han YC, et al. In vivo engineering of oncogenic chromosomal rearrangements with the CRISPR/Cas9 system. Nature 2014;516(7531):423–7 doi 10.1038/nature13902.

18. Yushkevich PA, Piven J, Hazlett HC, Smith RG, Ho S, Gee JC, et al. User-guided 3D active contour segmentation of anatomical structures: significantly improved efficiency and reliability. Neuroimage 2006;31(3):1116–28 doi 10.1016/j.neuroimage.2006.01.015.

19. Poczobutt JM, D. S, Yadav VK, Nguyen TT, Li H, Sippel TR, et al. Expression Profiling of Macrophages Reveals Multiple Populations with Distinct Biological Roles in an Immunocompetent Orthotopic Model of Lung Cancer. J Immunol 2016;196(6):2847–59 doi 10.4049/jimmunol.1502364.

20. Robinson MD, McCarthy DJ, Smyth GK. edgeR: a Bioconductor package for differential expression analysis of digital gene expression data. Bioinformatics 2010;26(1):139–40 doi 10.1093/bioinformatics/btp616.

21. Johnson AM, Boland JM, Wrobel J, Klezcko EK, Weiser-Evans M, Hopp K, et al. Cancer cell-specific MHCII expression as a determinant of the immune infiltrate organization and function in the non-small cell lung cancer tumor microenvironment. J Thorac Oncol 2021 doi 10.1016/j.jtho.2021.05.004.

22. Maynard A, McCoach CE, Rotow JK, Harris L, Haderk F, Kerr DL, et al. Therapy-Induced Evolution of Human Lung Cancer Revealed by Single-Cell RNA Sequencing. Cell 2020;182(5):1232–51 e22 doi 10.1016/j.cell.2020.07.017.

23. Kimball AK, Oko LM, Bullock BL, Nemenoff RA, van Dyk LF, Clambey ET. A Beginner’s Guide to Analyzing and Visualizing Mass Cytometry Data. J Immunol 2018;200(1):3–22 doi 10.4049/jimmunol.1701494.

24. Wang B, Krall EB, Aguirre AJ, Kim M, Widlund HR, Doshi MB, et al. ATXN1L, CIC, and ETS Transcription Factors Modulate Sensitivity to MAPK Pathway Inhibition. Cell reports 2017;18(6):1543–57 doi 10.1016/j.celrep.2017.01.031.

25. Xie Y, Cao Z, Wong EW, Guan Y, Ma W, Zhang JQ, et al. COP1/DET1/ETS axis regulates ERK transcriptome and sensitivity to MAPK inhibitors. J Clin Invest 2018;128(4):1442–57 doi 10.1172/JCI94840.

26. Johnson AM, Kleczko EK, Nemenoff RA. Eicosanoids in Cancer: New Roles in Immunoregulation. Frontiers in pharmacology 2020;11:595498 doi 10.3389/fphar.2020.595498.

27. Weiser-Evans MC, Wang XQ, Amin J, Van Putten V, Choudhary R, Winn RA, et al. Depletion of cytosolic phospholipase A2 in bone marrow-derived macrophages protects against lung cancer progression and metastasis. Cancer Res 2009;69(5):1733–8 doi 10.1158/0008-5472.CAN-08-3766.

28. Korpela SP, Hinz TK, Oweida A, Kim J, Calhoun J, Ferris R, et al. Role of epidermal growth factor receptor inhibitor-induced interferon pathway signaling in the head and neck squamous cell carcinoma therapeutic response. J Transl Med 2021;19(1):43 doi 10.1186/s12967-021-02706-8.

29. Fedele C, Li S, Teng KW, Foster CJR, Peng D, Ran H, et al. SHP2 inhibition diminishes KRASG12C cycling and promotes tumor microenvironment remodeling. J Exp Med 2021;218(1) doi 10.1084/jem.20201414.

30. Petroni G, Buque A, Zitvogel L, Kroemer G, Galluzzi L. Immunomodulation by targeted anticancer agents. Cancer Cell 2021;39(3):310–45 doi 10.1016/j.ccell.2020.11.009.

31. Budczies J, Kirchner M, Kluck K, Kazdal D, Glade J, Allgauer M, et al. Deciphering the immunosuppressive tumor microenvironment in ALK- and EGFR-positive lung adenocarcinoma. Cancer Immunol Immunother 2022;71(2):251–65 doi 10.1007/s00262-021-02981-w.

32. Ruscetti M, Leibold J, Bott MJ, Fennell M, Kulick A, Salgado NR, et al. NK cell-mediated cytotoxicity contributes to tumor control by a cytostatic drug combination. Science 2018;362(6421):1416–22 doi 10.1126/science.aas9090.

33. Canon J, Rex K, Saiki AY, Mohr C, Cooke K, Bagal D, et al. The clinical KRAS(G12C) inhibitor AMG 510 drives anti-tumour immunity. Nature 2019;575(7781):217–23 doi 10.1038/s41586-019-1694-1.

34. Tang KH, Li S, Khodadadi-Jamayran A, Jen J, Han H, Guidry K, et al. Combined Inhibition of SHP2 and CXCR1/2 Promotes Antitumor T-cell Response in NSCLC. Cancer discovery 2022;12(1):47–61 doi 10.1158/2159-8290.CD-21-0369.

35. Schreiber RD, Old LJ, Smyth MJ. Cancer immunoediting: integrating immunity’s roles in cancer suppression and promotion. Science 2011;331(6024):1565–70 doi 10.1126/science.1203486.

36. Liu SY, Dong ZY, Wu SP, Xie Z, Yan LX, Li YF, et al. Clinical relevance of PD-L1 expression and CD8+ T cells infiltration in patients with EGFR-mutated and ALK-rearranged lung cancer. Lung Cancer 2018;125:86–92 doi 10.1016/j.lungcan.2018.09.010.

