## Supplemental Figures 1-4 for "Adaptive immunity is required for durable responses to alectinib in murine models of EML4-ALK lung cancer"

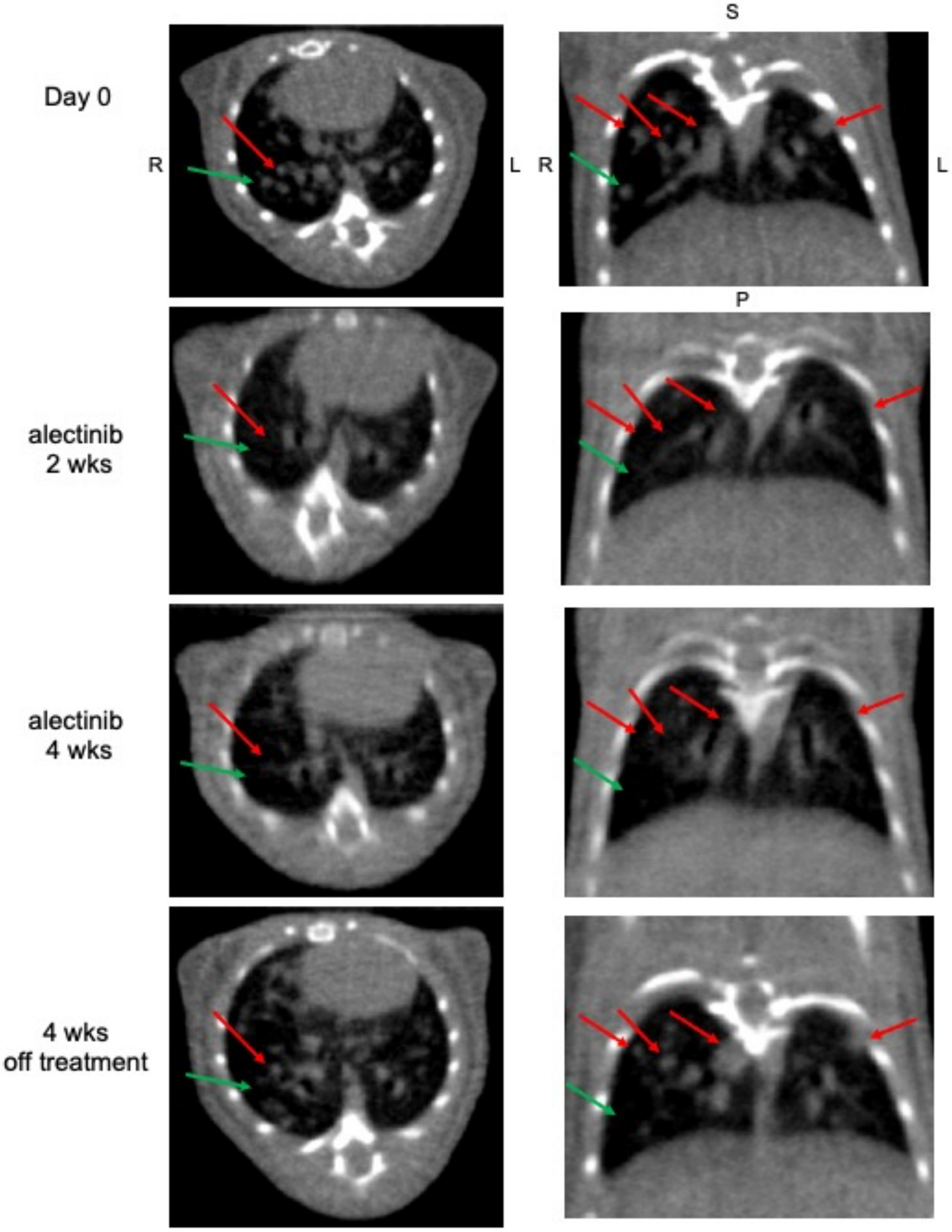


**Supplementary Figure S1. Alectinib sensitivity of primary murine EML4-ALK tumors.** Recombinant adenoviruses encoding Cas9 and gRNAs targeting *Eml4* and *Alk* were instilled intratracheally into C57BL/6 mice. The mice were routinely monitored by $\mu$CT for emergence of lung tumors and after 8 weeks (Day 0 in figure), mice were submitted to daily oral gavage with alectinib (20 mg/kg) and $\mu$CT continued on a weekly basis. Following 4 weeks of alectinib treatment, therapy was terminated and μCT imaging continued. Serial images of an alectinib-treated mouse (representative of 4 others) are shown. R = right, L = left, S = superior, P = posterior. Red arrows identify lesions that shrank upon alectinib treatment and subsequently re-grew following therapy termination. Green arrows identify lesions that shrank with treatment and did not regrow upon termination of alectinib therapy.

**A**

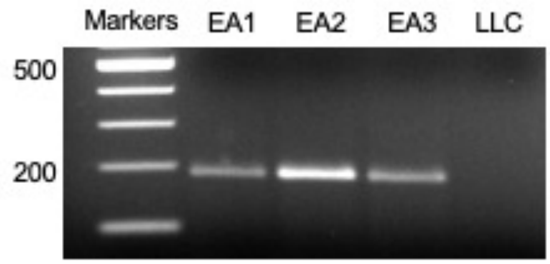


**B**

| RNA expression, CPM | | | | |
|---|---|---|---|---|
| Gene | EA1 | EA2 | EA3 | LLC |
| ALK | 646 | 1204 | 700 | 0 |
| MAPK1 | 1756 | 2602 | 1577 | 1664 |

**C**

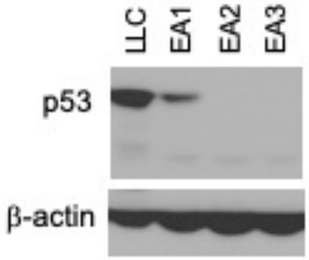


**Supplementary Figure S2**

**Supplementary Figure S2. RNA expression of EML4-ALK in murine cell lines. A,** RNA was purified from EA1, EA2 and EA3 cells and submitted to reverse transcription PCR to amplify the coding sequences expressed from the rearranged EML4-ALK gene fusion. **B**, RNAseq data from the cell lines was queried for ALK mRNA expression with MAPK1 as a control house-keeping gene. The LLC cell line is a KRAS mutant line that does not express ALK. **C**, Cell extracts were prepared from the indicated murine cell lines and submitted to immunoblot analysis for TP53 and β-actin as a loading control. LLC cells bear a missense mutation in TP53, EA1 was developed in TP53 wild-type mice and EA2 and EA3 were generated in TP53 null mice.

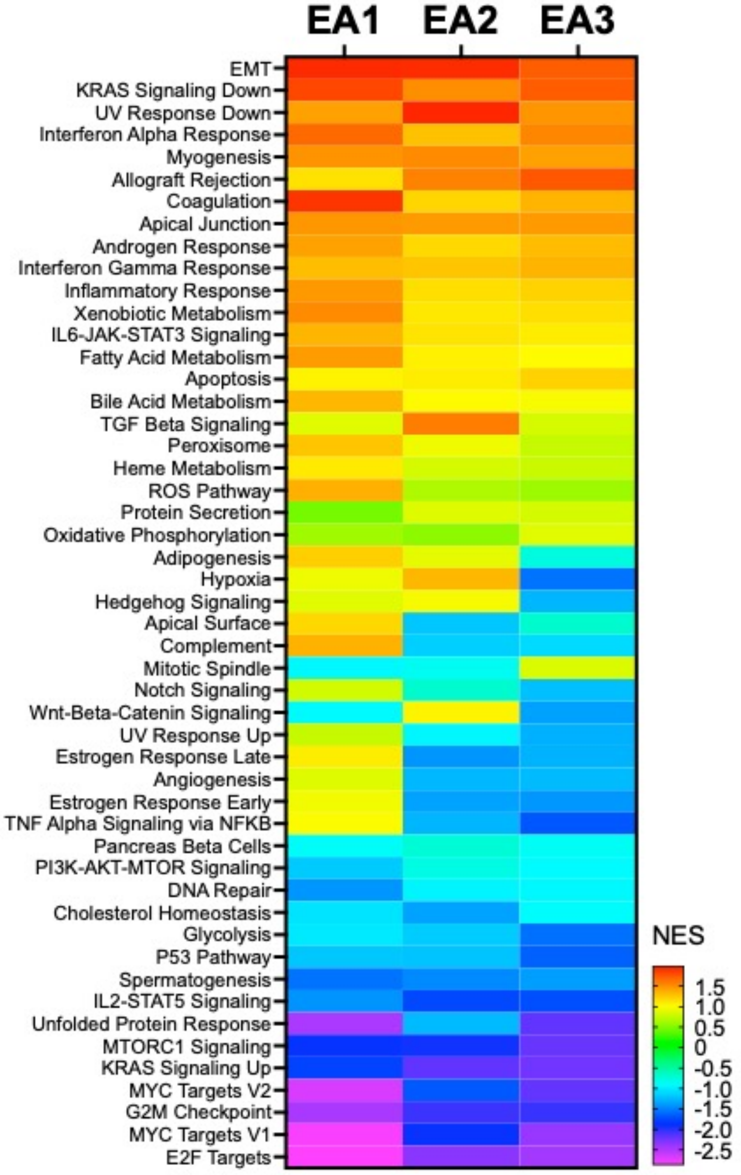


**Supplementary Figure S3**

**Supplementary Figure S3. Gene-set enrichment analysis of RNAseq data from murine EML4-ALK cell lines treated *in vitro* with alectinib.** RNA was purified from EA1, EA2 and EA3 cells treated *in vitro* with DMSO or alectinib (100 nM) for 1-5 days and submitted for RNAseq. For this analysis, the different time points were considered as replicates (n=4) and DMSO vs. alectinib-treated samples analyzed with

GSEA using the Hallmark Pathways. The heatmap presents the normalized enrichment scores (NES) in the alectinib-treated samples (see color bar for relative scores).

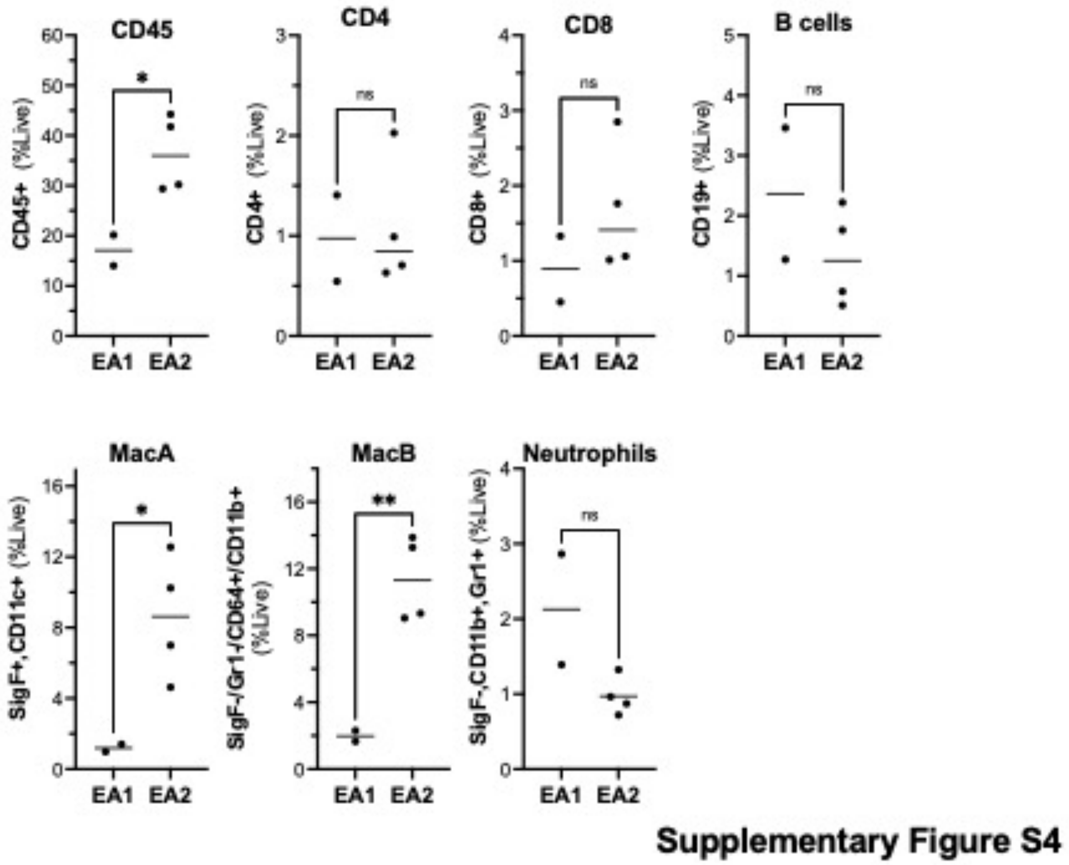


**Supplementary Figure S4. Baseline immune cell content in EA1 and EA2 orthotopic tumors.** EA1 or EA2 cells were implanted in the left lung of C57BL/6 mice and tumors were permitted to establish for 3 weeks. Left tumor-bearing lung lobes were dissociated, single cell suspensions were stained with 36 heavy metal-labelled antibodies, and samples ran on the Helios Mass Cytometer. The raw CyTOF data were de-barcoded, normalized and the live cells gated for further analyses. CD4+ T cells are CD45+/CD3+/CD4+. All CD8+ T cells are identified as CD45+/CD3+/CD8+. MacA, MacB, and neutrophil populations were gated as previously described in our flow cytometry (1). MacA cells correspond to resident lung macrophages and MacB cells are infiltrating macrophages. The different immune cell subtypes are presented as %Live and are derived from 2 (EA1) and 4 (EA2) independent tumor preparations. Comparisons were analyzed by Student's t-test where * and ** indicate p-values less than 0.05 and 0.01 respectively.


1. Poczobutt JM, De S, Yadav VK, Nguyen TT, Li H, Sippel TR*, et al.* Expression Profiling of Macrophages Reveals Multiple Populations with Distinct Biological Roles in an Immunocompetent Orthotopic Model of Lung Cancer. *J Immunol* 2016;**196**(6):2847-59 doi 10.4049/jimmunol.1502364.
